# A personal, reference quality, fully annotated genome from a Saudi individual

**DOI:** 10.1101/2022.11.05.515129

**Authors:** Maxat Kulmanov, Rund Tawfiq, Hatoon Al Ali, Marwa Abdelhakim, Mohammed Alarawi, Hind Aldakhil, Dana Alhattab, Ebtehal A. Alsolme, Azza Althagafi, Angel Angelov, Salim Bougouffa, Patrick Driguez, Yang Liu, Changsook Park, Alexander Putra, Ana M. Reyes-Ramos, Charlotte A. E. Hauser, Ming Sin Cheung, Malak S Abedalthagafi, Robert Hoehndorf

**Affiliations:** Computational Bioscience Research Center (CBRC), King Abdullah University of Science and Technology, Thuwal, Saudi Arabia; Computer, Electrical and Mathematical Sciences & Engineering (CEMSE) Division, King Abdullah University of Science and Technology, Thuwal, Saudi Arabia; Biological and Environmental Sciences & Engineering (BESE) Division, King Abdullah University of Science and Technology, Thuwal, Saudi Arabia; Laboratory for Nanomedicine, Biological and Environmental Science & Engineering (BESE) Division, King Abdullah University of Science and Technology (KAUST), Thuwal, Saudi Arabia; Genome Research Department, King Fahad Medical City, Riyadh, Saudi Arabia; Computer Science Department, College of Computers and Information Technology, Taif University, Taif, Saudi Arabia; Core Labs, King Abdullah University of Science and Technology, Thuwal, Saudi Arabia

## Abstract

We have used multiple sequencing approaches to sequence the genome of a volunteer from Saudi Arabia. We use the resulting data to generate a *de novo* assembly of the genome, and use different computational approaches to refine the assembly. As a consequence, we provide a contiguous assembly of the complete genome of an individual from Saudi Arabia for all chromosomes except chromosome Y, and label this assembly `KSA001`. We transferred genome annotations from reference genomes and predicted genome features using methods from Artificial Intelligence to fully annotate `KSA001`, and we make all primary sequencing data, the assembly, and the genome annotations freely available in public databases using the FAIR data principles.

## Introduction

The first complete, or almost complete, sequence of a human genome was made available in 2020 and published in 2022 [1], based on the functionally haploid cell line `CHM13`. Since then, several human genome assemblies were published [2], including the diploid genome sequences of 47 individuals [3]. The availability of these genomes is driven both by advances in sequencing technology which makes it possible to sequence more accurate and longer reads, and advances in assembly and read mapping algorithms [4–7] which can efficiently assemble and map sequence reads obtained from different sequencing technologies and of different quality, as well as assemble complex regions of genomes.

The availability of cheap, fast, and accurate sequencing technologies now enables sequencing of multiple genomes from diverse populations in order to understand their genetic variability. For each population, it also becomes possible to develop computational resources and databases that capture their diversity. Creating population-specific computational resources can serve as the foundation for bioinformatics workflows and thereby improve the accuracy of genomic analyses within a population as well as when comparing results between multiple different populations.

Reference genomes in particular are a foundation of workflows that analyze genomic data, and it is well-known that current reference genomes exhibit population bias [8] that may affect analyses built on them [9]. In the Middle East, the Qatari genome project [10] and the United Arab Emirates population genome project [11] aimed to address this challenge by producing a population-specific reference genome that includes as major alleles the most frequent ones identified within their respective populations.

However, many populations in the Middle East have historically been organized in tribal structures where marriages occur predominantly within a tribe [12] leading to several populations that were isolated for a period of time and therefore exhibit different genetic structure [13]. Additionally, the tribal structure also resulted in a relatively high prevalence of consanguineous marriages and consequently homozygosity and Mendelian disease [14]. No linear reference genome can fully capture the genetic diversity found in different populations within the Middle East.

To account for different populations within reference genomes, graph-based representations of multiple genomes are increasingly being used [15, 16]. In these genome (or pangenome) graphs, genomes from multiple individuals are represented and are used jointly for the alignment steps during bioinformatics workflows [17]. The Human Pangenome Project aims to generate a pangenome graph that captures the diversity across all human populations [18], with the initial effort focusing on genomic data derived from the 1,000 Genomes project which does not represent individuals from the Middle East well.

Although there are many genetic studies of ancient and current populations of the Middle East [19–22], only little genomic data from the Middle East is publicly available, or data that is available can only be used under prohibitive licenses and not be shared publicly. However, public availability and permissive licences are crucial for resources that need to be shared and utilized broadly, in particular for reference genomes. To further ensure broad usability, resources that underlie common analyses and workflows should be Findable, Accessible, Interoperable, and Reusable (FAIR) [23].

We sequenced the genome of a volunteer from Saudi Arabia who consented to make her genomic data public and freely available. We used three different sequencing strategies based on long and short reads. The first draft genome was created based on a *de novo* genome assembly using these reads. The assembly was further refined using the `CHM13` genome [1], resulting in the most complete publicly available genome sequence of a Saudi individual available so far. We use a variety of tools to fully annotate this genome, either transferring information from public databases or using methods from bioinformatics and Artificial Intelligence to identify functional element (genes, regulatory regions) using the genome sequence directly. Our sequencing, assembly, and annotation effort resulted in a reference-quality personal genome from a Saudi individual which we label `KSA001`. The assembled genome sequence of `KSA001`, the primary sequencing data, and the workflows used to construct the genome sequence are freely available on https://github.com/bio-ontology-research-group/KSA001. The sequence reads and assembly are further available in public sequence databases, and the genome is available in standard formats, making this one of the first FAIR individual genomes from the Arabian peninsula.

## Materials and Methods

### DNA extraction

DNA was extracted using two methods. Ultra High molecular weight DNA (uHMW) was isolated from fresh blood samples using New England Biolabs (NEB) Monarch High Molecular Weight (HMW) DNA isolation kit following manufacturer protocol (New England Biolabs, UK) with modification, agitation was set at 700 rpm during the lysis step. DNA was kept at 4°C until library preparation for long-read sequencing using the PacBio andOxford nanopore sequencing platforms.

For short reads sequencing, We isolated donor DNA using the DNeasy Blood & Tissue kit (Qiagen) following the manufacturers instructions. DNA was kept at -20°C until library preparation.

### Sequencing library preparation and sequencing

We use three platforms for sequencing the extracted DNA: Illumina NovaSeq 6000, Pacbio Sequel II, and Oxford Nanopore PromethION.

We used 100ng of genomic DNA (gDNA) as an input to construct a whole genome library to be sequenced using the NovaSeq 6000 platform. The DNA was mechanically sheared with Covaris and converted to a sequence-ready library using the TruSeq DNA Nano Library Kit (Illumina, USA). Subsequently, the library was quantified using Qubit high-sensitivity dsDNA Assays (ThermoFisher Scientific Q33230, USA), and the quality control was performed using an Agilent Bioanalyzer 2100 (Agilent Technologies, USA). The sample was sequenced on two lanes of an SP flowcell with a read length of 2 *×* 150 bp in paired-end format, which generated 311 GB of data, resulting in an estimated 92x coverage.

High molecular weight gDNA was sheared with Megaruptor 3 (Diagenode, Denville, USA) to the size range of 15-20kb. SMRTbell was prepared with HiFi Express Template prep kit 2.0 (102-088-900), and size-selected with the PippinHT System (Sage Science HTP0001). Finally, SMRTbell QC was assessed with Qubit dsDNA High Sensitivity (ThermoFisher Scientific Q33230) and FEMTO Pulse (Agilent Technologies, Inc. P-0003-0817). Sequencing of SMRTbell was set up on PacBio Sequel II system with Sequel II Binding kit 2.2 (101-894-200), Sequel II Sequencing Kit 2.0 (101-820-200), and SMRT cell 8M Tray (101-389-001), according to conditions specified in SMRTlink with 30 hour movie times, 2 hour pre-extension time, and adaptive loading mode.

For Nanopore sequencing, two libraries were prepared using the Ultra-Long Sequencing Kit (SQK-ULK001) and its recommended protocol from Oxford Nanopore Technologies (ONT). 30 µg of uHMW gDNA was used as input for each library. A PromethION flow cell (FLO-PRO002) was primed and prepared according to the same protocol and 75 µl of a sequencing library was loaded. 24 hours after the first sequencing run, a nuclease flush and priming step were performed according to the protocol, and an additional 75 µl of library was loaded before starting another sequencing run. This process was repeated until each library was loaded three times. Basecalling was performed using Guppy v5.1.13 with the super-accurate basecalling model. The entire process is repeated with the second prepared library and another PromethION flow cell.

### Draft assembly

We used PacBio HiFi reads resulting from Circular Consensus Sequencing (CCS) and independently applied Hifiasm v0.16.1 [24] and Flye v2.9.1 [25] for the assembly. The ONT reads were also assembled using Flye v2.9.1. Our best assembly was generated by Hifiasm from HiFi reads and resulted in 159 contigs. Table 1 provides statistics on each of our assemblies. We use the Hifiasm assembly as our primary assembly and use the Flye assemblies to bridge the contigs and fill the gaps between them.

**Table 1.** Assembly statistics

| Tool | Contigs | N/L50 | N/L90 | Length |
| --- | --- | --- | --- | --- |
| Hifiasm | 159 | 11/105.778 MB | 17/80.051 MB | 3063.335 MB |
| Flye Hifi | 27,170 | 628/1.287 MB | 885/908.773 KB | 3734.855 MB |
| Flye Ont | 780 | 19/55.843 MB | 35/32.797 MB | 2860.535 MB |

After initial HiFi assembly, we used the chromosome scaffolder from MaSuRCA v4.1.0 [26] to break misassemblies and place contigs in the right location within their respective chromosomes. We used `CHM13` [1] as a reference and PacBio HiFi reads as input raw reads. The tool aligns the assembly to the reference and identifies alignment breakpoints. The HiFi reads are then aligned to the assembly and coverage around the alignment breakpoints is examined. Very low or very high coverage around the breakpoints indicates a possible misassembly. Flagged misassemblies are broken and re-aligned against the reference for the final re-assembly. This step produced 104 scaffolds and 276 contigs.

Furthermore, we used the SAMBA scaffolder [27] tool to fill in the gaps and connect the contigs. The SAMBA scaffolder uses additional long read data to increase the contiguity of the assemblies. We used both HiFi and ONT reads, and the Flye generated assemblies with minimum match length on both sides of the gap of 2500. In total, we filled 60 gaps in this step. The resulting genome (v0.2.0) contained 90 scaffolds and 216 contigs.

### Refinement of assembly

After this initial assembly, we aligned the contigs to `CHM13` using minimap2 [28] and manually inspected the gaps that were not linked in previous steps. We found that most of them are in centromeres which contain highly repetitive genome and are generally difficult to align and assemble [29, 30]. Based on this finding, we decided to assemble centromeric regions separately. We removed the centromeric regions from `CHM13` and aligned our contigs again. By placing our contigs on the reference genome we were able to close all the gaps in non-centromeric regions except approximately a 2 MB region in chromosomes 16 and X, and the entire mitochondrial genome.

We aligned the contigs in our ONT-based assembly to `CHM13` in order to inspect if they include missing parts from the main assembly. We found that the gaps in chromosomes 16 and X, and the complete mitochondrial genome, were assembled in our ONT assembly. Therefore, we merged them with our main assembly.

For centromeric regions, we found that in chromosomes 2, 5, 7, 8, 10, 17, 18, 19 and 20 we have a single contig and in chromosomes 1 and 3 three contigs which cover the entire region. The centromeric regions in chromosomes 4, 6, 9, 11, 12, 13, 14, 15, 16, 21, 22, and X, were not assembled well and fragmented into multiple contigs that did align well to `CHM13`. For these regions we partially copied the missing parts from `CHM13`.

Furthermore, we polished regions admitted from the ONT-based assembly using NextPolish [31] guided by quality-filtered Illumina reads to reduce base errors of ONT assembled sequences.

We calculated QV scores using Merqury [32] with a *k*-mer size of 25.

### Genome annotations

We annotated the `KSA001` genome based on Gencode (CAT-Liftoff) annotation of `CHM13` v2.0 [1] using Liftoff v1.6.1 [33] with the following parameters: `-sc 0.95 –polish -chroms -exclude partial -copies`. We then reapplied LiftOff with the option `-overlap 1.0`.

We performed *ab initio* gene prediction using Augustus v3.3.3 [34] with the following parameters `--strand=both --genemodel=complete --singlestrand=true --gff3=on --softmasking=on --species=human`. To mask repeats in the FASTA file, we used RepeatMasker v4.1.1 [35] with the parameters `-species human -xsmall`. We used gffcompare [36] v0.12.6 to compare the *de novo* gene annotation with remapped genes.

Prediction of enhancers, transcription start sites (TSS), histone modification marks and gene expression was done using Enformer [37] on both the `KSA001` and `CHM13` genomes. Scores for all generated tracks were converted to Wig files and compared to Encode annotations converted to `KSA001` using CrossMap v0.6.4.

To use `KSA001` in downstream applications, we generated chain files between `KSA001` (v0.1.0) and both `CHM13` (v2.0) and `GRCh38` (v41) using pyOverChain. We then applied LiftOver on the entire dbSNP (v155) database from `GRCh38` to `KSA001` (v0.1.0) using LiftoverVcf from Picard (v2.20.4).

### Variant calling

We performed two types of variant calling. First, to analyze variant differences, we aligned `KSA001` to `CHM13` using Minimap2 (v2.24-r1122) and used the call command within pftools.js to call variants. Results were filtered to remove variants in centromeric regions as we copied some centromeric regions from `CHM13` into `KSA001`. A Harr plot of the assembly was generated using minidot from miniasm (v0.3-r179) [38].

A second type of variant calling was performed to compare `KSA001` against `CHM13` as a reference genome for performing variant calling on Saudi individuals. We utilized an in-house cohort of Saudi genomes, sequenced on a NovaSeq 6000 S4 flow cell with 150 *×* 150 bp read lengths, an average genome-wide coverage of 30x, and sequencing depth around 50x-60x. The cohort consisted of individuals diagnosed with hepatocellular carcinoma (HCC) with DNA isolated from both tumors and blood samples. We used the germline genomes of 6 individuals.

To analyze variant differences, we aligned Illumina reads of each individual genome to `KSA001`, `GRCh38`, and `CHM13` using BWA-MEM (v0.7.17) [39]. We then sorted and indexed the alignment files using Picard (v2.20.4), and marked duplicate reads using the same tool. To reduce, systematic errors in base quality scores caused by sequencers, we performed base quality score reacalibration (BQSR) using GATK (v4.1.2.0) by first building a covariation model and then applying it to adjust the quality scores based on the model. For variant calling, we used HaplotypeCaller within GATK to call SNPs and small indels through a local *de novo* assembly of haplotypes in specific regions.

### Ethical approval

This work was approved by the Institutional Review Board (IRB) at the Faculty of Medicine, King Fahad Medical City, under approval number `22-037`, and by the Institutional Bioethics Committee (IBEC) at King Abdullah University of Science and Technology under approval number `22IBEC023`. The volunteer donor provided an informed consent to make the data resulting from genome sequencing of the donated DNA public. Use of the HCC genetic data was approved by the IRB at the Faculty of Medicine, King Fahad Medical City, under approval number `19-297`, and by the Institutional Bioethics Committee (IBEC) at King Abdullah University of Science and Technology under approval number `19IBEC29`.

### Data availability

Data, supporting information, and a description of the assembly workflow are available at https://github.com/bio-ontology-research-group/KSA001. Sequencing reads are available on the Sequence Read Archive under accession numbers SRR21927836, SRR21927835, SRR21927834, and SRR21927833.

## Results

### KSA001, and comparison to CHM13 and GRCh38

We generated the complete assembly of a Saudi individual for non-centromeric regions, and a partially complete assembly for centromeric regions. Although we sequence a diploid genome, due to the lack of long-range information from sequencing and lack of parental genotype data, we do not provide a diploid assembly but rather focus on generating a reference-quality assembly with high quality annotations.

The first version of the assembly (`KSA001` v0.1.0) was generated using only PacBio HiFi reads using the Hifiasm assembler [24]; this first assembly contained 137 contigs. After generating scaffolds by aligning to the `CHM13` genome [1], we found that a large part of chromosome 16 and the entire mitochondrial genome were missing. We then used Hifiasm without any post-joining steps to generate another assembly (v0.2.0) which included 159 contigs. After aligning to `CHM13`, this assembly covered most of all chromosomes except the mitochondrial genome and around 2 MBs in chromosomes 16 and X. We filled these gaps using an assembly based on ONT long reads, generated using the Flye assembler [25], to generate version v0.2.1 of `KSA001`. Version v0.3.0 of `KSA001` is based on applying NextPolish to polish the regions copied from the ONT based assembly in order to generate an assembly that is accurate on a basepair level. In version v0.3.0, centromeric regions in chromosomes 4, 6, 9, 11, 12, 13, 14, 15, 16, 21, 22, and X, were not assembled well and split into multiple contigs which did not align to centromeric regions in `CHM13`. We filled the gaps by copying them from `CHM13`. In total, we copied 114, 245, 615 bases from `CHM13` centromeres (3.7% of the total assembly).

Table 2 shows the completeness of the chromosomes and the number of basepairs in each chromosome of `KSA001` in comparison to the `CHM13` v2.0 assembly. Similar to `CHM13`, `KSA001` contains a single contig per chromosome, and the overall number of base-pairs in each chromosome are comparable to `CHM13`. Also, we report base-level quality (QV) scores for the `KSA001` assembly for each chromosome and the entire assembly. QV score represents a log-scaled probability of error for the consensus base calls.

**Table 2.**
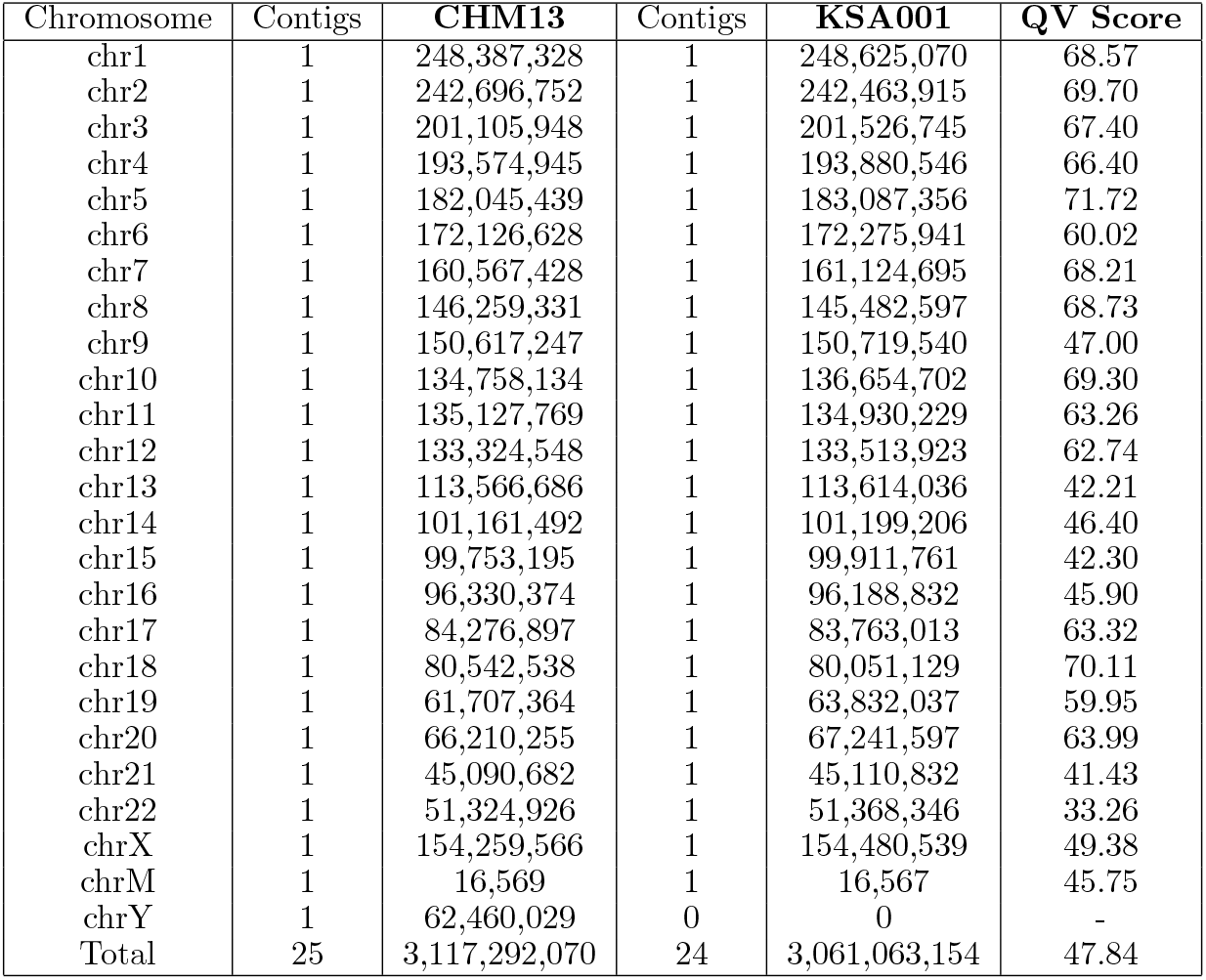
Completeness of chromosomes of `KSA001` compared to `CHM13` and QV scores.

To further compare `KSA001` and `CHM13`, we aligned the assemblies of `KSA001` (v0.3.0) and `CHM13` (v2.0) and counted the number and type of differences (SNPs, indels, structural variants). Table 3 shows the results of this comparison. The Harr plot (Figure 1) shows co-linearity of the two genomes with a region of inversion in chromosome 8.

**Table 3.**
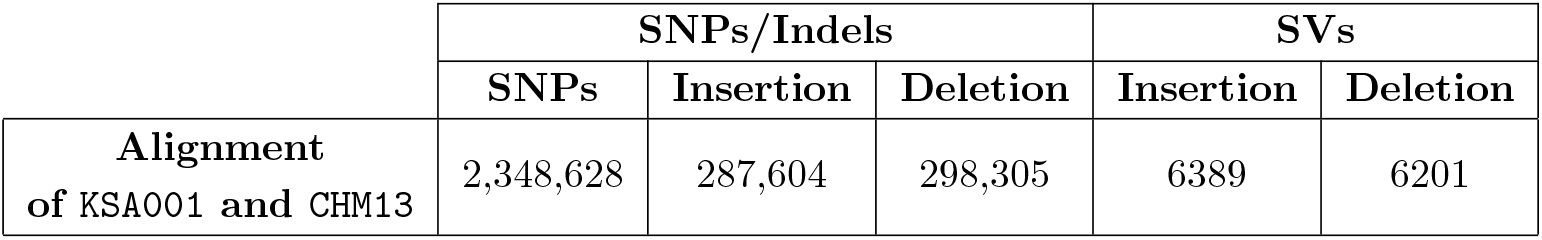
Comparison of `KSA001` and `CHM13` based on assembly-to-assembly alignment. SNPs: Single nucleotide polymorphisms, Indels: insertion or deletion, SV: Structural Variants

**Figure 1.**
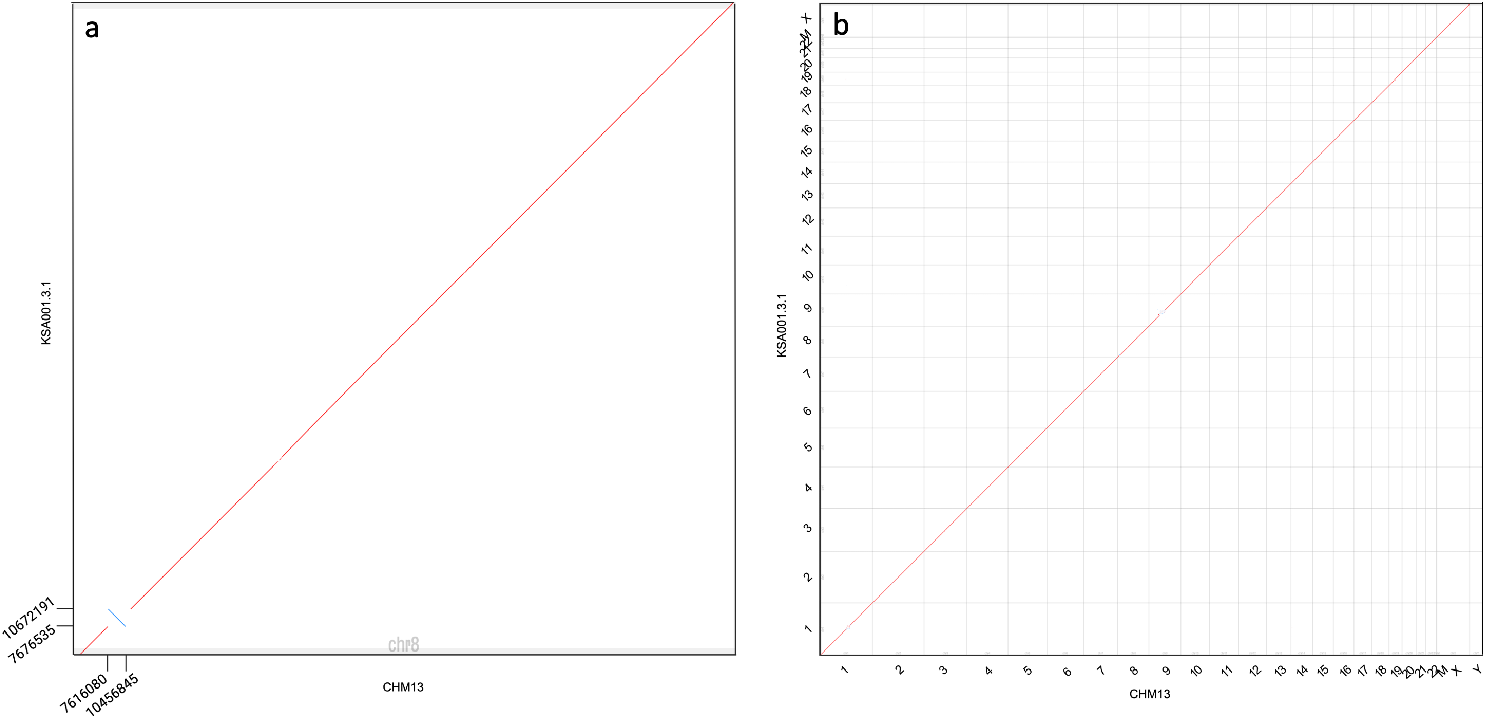
Harr plot of `KSA001` aligned with `CHM13`. (a) Plot of chromosome 8 in both genomes showing an inversion; b) plot of the entire alignment.

### Using KSA001 in variant calling workflows

In addition to the comparison of the two assemblies `KSA001` and `CHM13`, we also perform variant calling using short read sequencing. First, we perform variant calling of short reads derived from sequencing `KSA001`, using `CHM13` and `GRCh38` as reference genomes, and report the number of variants in `KSA001` when aligned against both the reference genomes (Table 4. Consistent with previously reported results [9], we observe a substantially lower number of variants called when using `CHM13` as a reference compared to using `GRCh38`.

**Table 4.** Variant calls statistics for `KSA001` Illumina reads aligned to `GRCh38` and `CHM13`

|  | Total Number of variants | SNPs | Indels |
| --- | --- | --- | --- |
| **KSA001 aligned to GRCh38** | 5,337,778 | 4,317,999 | 1,115,246 |
| **KSA001 aligned to CHM13** | 4,888,875 | 3,934,939 | 1,036,797 |

Saudi Arabia, and the MENA region, have populations with a high degree of consanguineous marriages, and accurate calling of genomic variants is particularly important for the diagnosis of Mendelian diseases. Furthermore, structural variants are more challenging to identify from sequencing data, and population-specific structural variants are not included in reference genomes. We have identified multiple structural variants (Table 3) ranging up to 140, 031 bp in size in `KSA001` when compared to `CHM13`, and including common structural variants in a reference genome used for variant calling can improve accuracy in determining the presence of variants and therefore the sensitivity in finding disease-causing variants.

The majority of genomic samples, in particular in a clinic, are still processed using short-read sequencing methods. To test whether `KSA001` can improve variant calling in Saudi individuals, we used 6 germline DNA samples extracted from Saudi individuals (HCC cohort), and performed variant calling of these samples using `CHM13` v2.0 and `KSA001` (v0.1.0) as references. As part of this process, we also generated a number of common resources that transfer information from `CHM13` to `KSA001`. Table 5 shows a summary of the average number of variants called on the HCC cohort we are using. We identify fewer average number of variants when using `KSA001` compared to using `CHM13`, indicating that `KSA001` captures major alleles in the Saudi population better than `CHM13`.

**Table 5.** Variant calling statistics for the HCC cohort, using Illumina reads aligned to `KSA001` and `CHM13`

|  | Total Number of variants | SNPs | Indels | Homozygous variants | Heterozygous variants |
| --- | --- | --- | --- | --- | --- |
| **KSA001** | 4,611,503 | 3,733,094 | 882,013 | 1,510,112 | 3,180,981 |
| **CHM13** | 5,011,016 | 4,080,430 | 934,495 | 1,632,735 | 3,453,533 |

### Genome annotation

We first annotated the `KSA001` genome by transferring genome features from `CHM13` to `KSA001` using the Liftoff tool [33]. Liftoff successfully mapped 62,864 genes out of 63,494 genes (99%, Table 6) to `KSA001`. When we further considered overlapping genome feature annotations, 520 additional genes were mapped. A gene is considered mapped when it satisfies at least 50% of alignment coverage and sequence identity. Most annotated genes were mapped with coverage and sequence identity higher than 99.5% (95% and 86% of genes, respectively). Out of the 110 unmapped genes (Supplementary Table 7), 38 were mapped partially. Supplementary Table 8 provides a summary of all mapped genome features.

**Table 6.** LiftOff annotation statistics. Comparison between numbers of genes and transcripts in `CHM13` and `KSA001`

|  | Genes |  | Transcripts |  |
| --- | --- | --- | --- | --- |
|  | Total number | Protein coding | Total number | Protein coding |
| **KSA001** | 63,384 | 19,956 | 233,476 | 155,931 |
| **CHM13** | 63,494 | 19,969 | 233,615 | 155,972 |

NTAN1 is the only protein-coding gene completely missing from `KSA001`; however, multiple pseudogenes orthologous to NTAN1 are present. Furthermore, there are 12 protein-coding genes partially mapped; two of these are olfactory receptors.

## Discussion

We sequenced a genome from an individual in Saudi Arabia, assembled and annotated it, and made it publicly available. The genome has been assembled to a very high level of completeness, comparable in completeness to the telomere-to-telomere assembled genome `CHM13`, with the exception that chromosome Y is entirely missing.

We make this genome publicly available following the FAIR principles [23]. The genome is findable as it is deposited in repositories containing genome assemblies, and accessible through common protocols and data download utilities. It is also interoperable as we use standard formats used across bioinformatics, and it is reusable as we make the primary data available as well as a description of the workflow that led to the assembled genome. The quality of the assembly and the set of genome annotations we make available, but in particular its free availability, allows `KSA001` to be shared and used as reference genome for genomic studies in Saudi Arabia and the Middle East. We demonstrated that `KSA001` can be used for variant calling and therefore contribute to the success of diagnostic or prognostic genomic methods.

In the future, we will extend the assembly to a diploid assembly, and we will update the `KSA001` assembly accordingly. A diploid assembly may become possible when more data becomes available, in particular, Hi-C sequencing data. Even more important than generating a diploid assembly, however, is sequencing and assembling more genomes from the Middle East and making the genome sequences available so that it becomes possible to construct a pangenome that captures the entire genetic diversity found within the Middle East. `KSA001`, as the first publicly available complete genome from a Saudi individual, can be used to start such an initiative.

## Acknowledgments

This work has been supported by funding from King Abdullah University of Science and Technology (KAUST) Office of Sponsored Research (OSR) under Award No. URF/1/4355-01-01, URF/1/4675-01-01, URF/1/4697-01-01, FCC/1/1976-46-01 and FCC/1/1976-34-01. MSA is supported by King Salman Center for disability research grant R-20190016. We acknowledge support from the KAUST Supercomputing Laboratory, and support from the KAUST Bioscience Core Laboratory.

## Author contributions

MK: draft assembly and refinement, analysis; RT: variant calling, analysis; HAA: genome annotation, inversions; MAlarawi: DNA extraction; MAbdelhakim: sample processing, DNA extraction, library preparation; HA: library preparation; EA: sample processing, DNA extraction; DA: DNA extraction; AAlth: variant calling, analysis; AAng: PacBio and ONT protocol optimization and sequencing; SB: draft assembly and refinement; YL: variant calling; PD: PacBio and ONT protocol optimization and sequencing, supervision; CP: PacBio and ONT protocol optimization and sequencing;AP: PacBio and ONT protocol optimization and sequencing; ARR: PacBio and ONT protocol optimization and sequencing; NC: supervision; MSA: conception, sample processing, supervision, funding acquisition; RH: conception, supervision, funding acquisition.

## Supplementary information

**Table 7.**
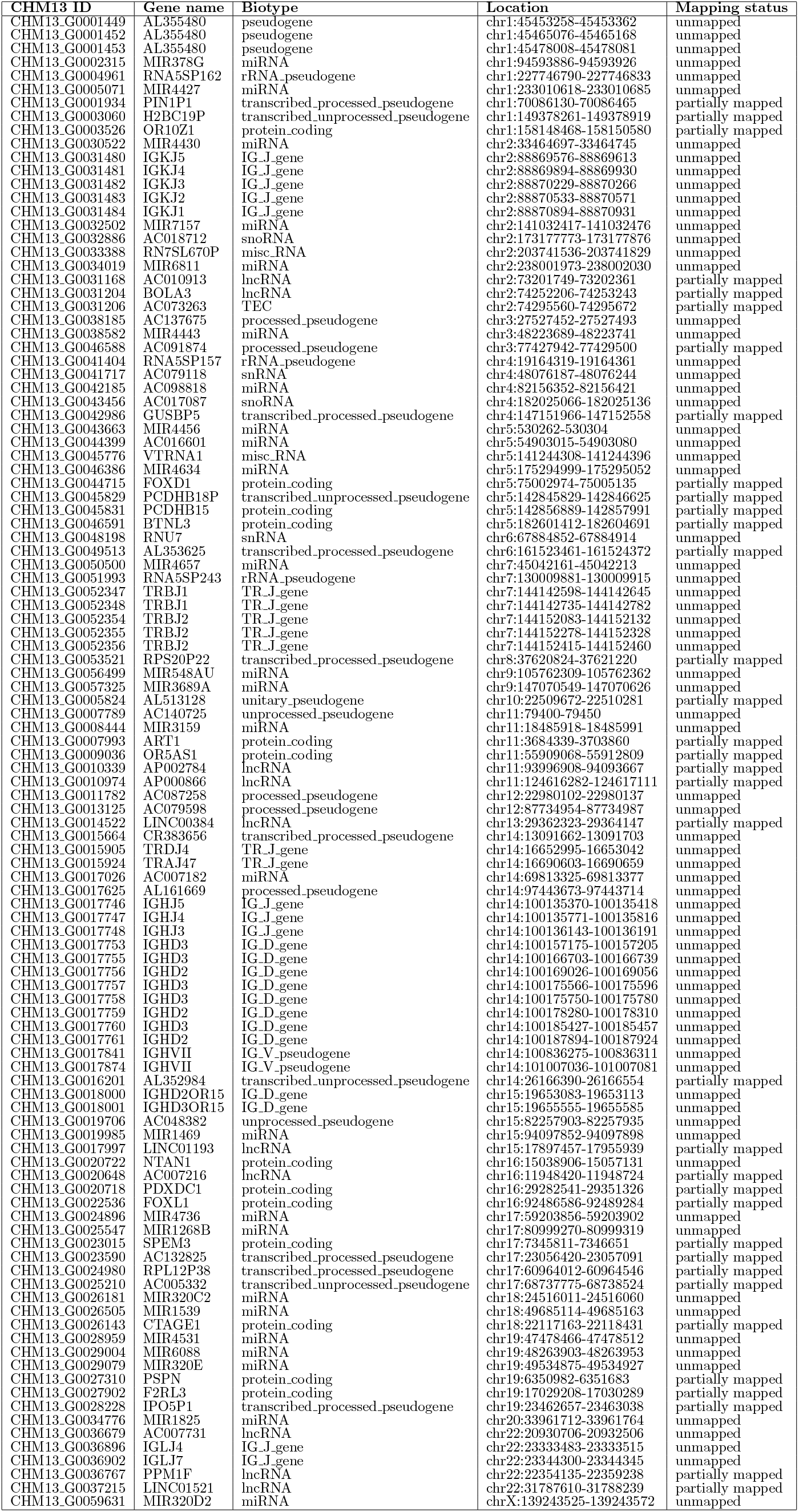
Unmapped geenes. List of completely missing and partially mapped genes (coverage and sequence identity *<* 50%)

**Table 8.**
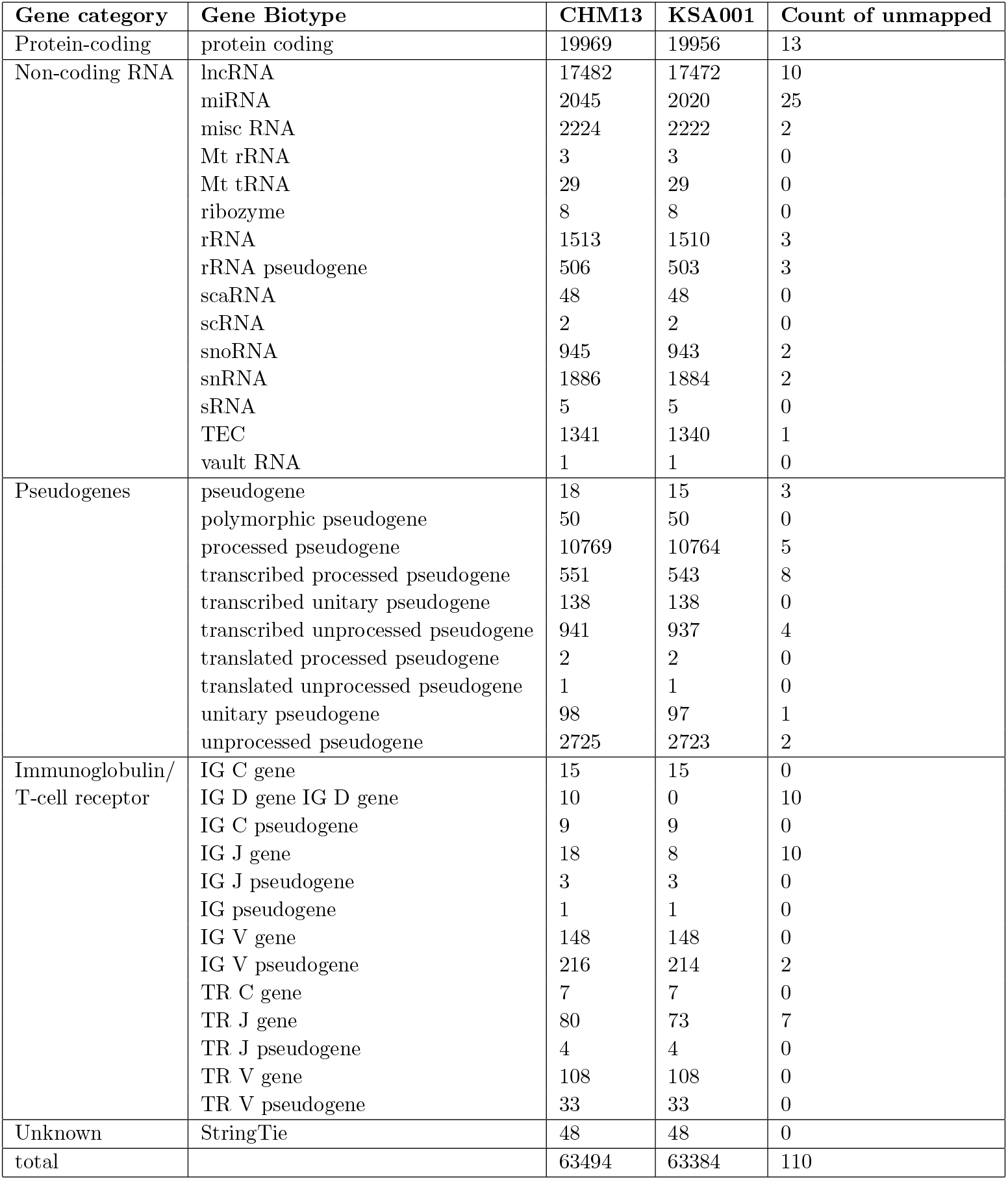
Gene annotation summary. Counts of `CHM13` genes mapped by Liftoff to the `KSA001` genome assembly.

## References

[1] S. Nurk et al. “The complete sequence of a human genome”. In: Science 376.6588 (2022), pp. 44–53.

[2] A. V. Zimin et al. “A reference-quality, fully annotated genome from a Puerto Rican individual”. In: Genetics 220.2 (2022), iyab227.

[3] W.-W. Liao et al. “A Draft Human Pangenome Reference”. In: bioRxiv (2022).

[4] A. M. Mc Cartney et al. “Chasing perfection: validation and polishing strategies for telomere-to-telomere genome assemblies”. en. In: Nature Methods 19.6 (2022), pp. 687–695.

[5] C. Jain, A. Rhie, N. F. Hansen, S. Koren, and A. M. Phillippy. “Long-read mapping to repetitive reference sequences using Winnowmap2”. en. In: Nature Methods 19.6 (2022), pp. 705–710.

[6] M. R. Vollger et al. “Improved assembly and variant detection of a haploid human genome using single-molecule, high-fidelity long reads”. en. In: Annals of Human Genetics 84.2 (2020), pp. 125–140.

[7] S. Nurk et al. “HiCanu: accurate assembly of segmental duplications, satellites, and allelic variants from high-fidelity long reads”. en. In: Genome Research 30.9 (2020), pp. 1291–1305.

[8] B. Paten, A. M. Novak, J. M. Eizenga, and E. Garrison. “Genome graphs and the evolution of genome inference”. en. In: Genome Research 27.5 (2017), pp. 665–676.

[9] S. Aganezov et al. “A complete reference genome improves analysis of human genetic variation”. In: Science 376.6588 (2022), eabl3533.

[10] K. A. Fakhro et al. “The Qatar genome: a population-specific tool for precision medicine in the Middle East”. en. In: Human Genome Variation 3.1 (2016), pp. 1–7.

[11] G. Daw Elbait, A. Henschel, G. K. Tay, and H. S. Al Safar. “A Population-Specific Major Allele Reference Genome From The United Arab Emirates Population”. In: Frontiers in Genetics 12 (2021).

[12] O. Bakoush, A. Bredan, and S. Denic. “KIN AND NON-KIN MARRIAGES AND FAMILY STRUCTURE IN A RICH TRIBAL SOCIETY”. en. In: Journal of Biosocial Science 48.6 (2016), pp. 797–805.

[13] K. Mineta, K. Goto, T. Gojobori, and F. S. Alkuraya. “Population structure of indigenous inhabitants of Arabia”. en. In: PLOS Genetics 17.1 (2021), e1009210.

[14] F. S. Alkuraya. “Genetics and genomic medicine in Saudi Arabia”. en. In: Molecular Genetics & Genomic Medicine 2.5 (2014), pp. 369–378.

[15] G. Hickey et al. “Genotyping structural variants in pangenome graphs using the vg toolkit”. In: Genome Biology 21.1 (2020), p. 35.

[16] J. M. Eizenga et al. “Pangenome Graphs”. In: Annual Review of Genomics and Human Genetics 21.1 (2020), pp. 139–162.

[17] D. Kim, J. M. Paggi, C. Park, C. Bennett, and S. L. Salzberg. “Graph-based genome alignment and genotyping with HISAT2 and HISAT-genotype”. en. In: Nature Biotechnology 37.8 (2019), pp. 907–915.

[18] T. Wang et al. “The Human Pangenome Project: a global resource to map genomic diversity”. en. In: Nature 604.7906 (2022), pp. 437–446.

[19] S. Mallick et al. “The Simons Genome Diversity Project: 300 genomes from 142 diverse populations”. en. In: Nature 538.7624 (2016), pp. 201–206.

[20] S. E. John et al. “Assessment of coding region variants in Kuwaiti population: implications for medical genetics and population genomics”. en. In: Scientific Reports 8.1 (2018), p. 16583.

[21] I. Lazaridis et al. “Genomic insights into the origin of farming in the ancient Near East”. en. In: Nature 536.7617 (2016), pp. 419–424.

[22] E. M. Scott et al. “Characterization of Greater Middle Eastern genetic variation for enhanced disease gene discovery”. en. In: Nature Genetics 48.9 (2016), pp. 1071–1076.

[23] M. D. Wilkinson et al. “The FAIR Guiding Principles for scientific data management and stewardship”. en. In: Scientific Data 3.1 (2016), p. 160018.

[24] H. Cheng, G. T. Concepcion, X. Feng, H. Zhang, and H. Li. “Haplotype-resolved de novo assembly using phased assembly graphs with hifiasm”. en. In: Nature Methods 18.2 (2021), pp. 170–175.

[25] Y. Lin et al. “Assembly of long error-prone reads using de Bruijn graphs”. In: Proceedings of the National Academy of Sciences 113.52 (2016), E8396–E8405.

[26] A. V. Zimin et al. “Hybrid assembly of the large and highly repetitive genome of Aegilops tauschii, a progenitor of bread wheat, with the MaSuRCA mega-reads algorithm”. In: Genome Research 27.5 (2017), pp. 787–792.

[27] A. V. Zimin and S. L. Salzberg. “The SAMBA tool uses long reads to improve the contiguity of genome assemblies”. en. In: PLOS Computational Biology 18.2 (2022), e1009860.

[28] H. Li. “New strategies to improve minimap2 alignment accuracy”. In: Bioinformatics 37.23 (2021), pp. 4572–4574.

[29] K. E. Hayden. “Human centromere genomics: now it’s personal”. In: Chromosome Research 20.5 (2012), pp. 621–633.

[30] N. Altemose et al. “Complete genomic and epigenetic maps of human centromeres”. In: Science 376.6588 (2022), eabl4178.

[31] J. Hu, J. Fan, Z. Sun, and S. Liu. “NextPolish: a fast and efficient genome polishing tool for long-read assembly”. In: Bioinformatics 36.7 (2020), pp. 2253–2255.

[32] A. Rhie, B. P. Walenz, S. Koren, and A. M. Phillippy. “Merqury: reference-free quality, completeness, and phasing assessment for genome assemblies”. In: Genome Biology 21.1 (2020), p. 245.

[33] A. Shumate and S. L. Salzberg. “Liftoff: accurate mapping of gene annotations”. In: Bioinformatics 37.12 (2021), pp. 1639–1643.

[34] O. Keller, M. Kollmar, M. Stanke, and S. Waack. “A novel hybrid gene prediction method employing protein multiple sequence alignments”. In: Bioinformatics 27.6 (2011), pp. 757–763.

[35] M. Tarailo-Graovac and N. Chen. “Using RepeatMasker to Identify Repetitive Elements in Genomic Sequences”. en. In: Current Protocols in Bioinformatics 25.1 (2009), pp. 4.10.1–4.10.14.

[36] G. Pertea and M. Pertea. “GFF utilities: GffRead and GffCompare”. In: F1000Research 9 (2020).

[37] Ž. Avsec et al. “Effective gene expression prediction from sequence by integrating long-range interactions”. In: Nature methods 18.10 (2021), pp. 1196–1203.

[38] H. Li. “Minimap and miniasm: fast mapping and de novo assembly for noisy long sequences”. In: Bioinformatics 32.14 (2016), pp. 2103–2110.

[39] H. Li. Aligning sequence reads, clone sequences and assembly contigs with BWA-MEM. 2013.

